# Activation of the SARS-CoV-2 receptor *Ace2* by cytokines through pan JAK-STAT enhancers

**DOI:** 10.1101/2020.05.11.089045

**Authors:** Lothar Hennighausen, Hye Kyung Lee

## Abstract

ACE2, in concert with the protease TMPRSS2, binds the novel coronavirus SARS-CoV-2 and facilitates its cellular entry. The *ACE2* gene is expressed in SARS-CoV-2 target cells, including Type II Pneumocytes (Ziegler, 2020), and is activated by interferons. Viral RNA was also detected in breast milk (Wu et al., 2020), raising the possibility that *ACE2* expression is under the control of cytokines through the JAK-STAT pathway. Here we show that *Ace2* expression in mammary tissue is induced during pregnancy and lactation, which coincides with the establishment of a candidate enhancer. The prolactin-activated transcription factor STAT5 binds to tandem sites that coincide with activating histone enhancer marks and additional transcription components. The presence of pan JAK-STAT components in mammary alveolar cells and in Type II Pneumocytes combined with the autoregulation of both STAT1 and STAT5 suggests a prominent role of cytokine signaling pathways in cells targeted by SARS-CoV-2.

## Introduction

ACE2, the receptor for SARS-CoV (Imai et al., 2005) and SARS-CoV-2 (Hoffmann et al., 2020), has been identified in several target cells, including absorptive Enterocytes (Lamers et al., 2020), secretory goblet cells (Zhao, 2020), the olfactory system (Brann, 2020) and several epithelial cell types (Brann, 2020; Lukassen et al., 2020; Qi et al., 2020). A study in pneumocytes demonstrated that *ACE2* expression is induced by interferons (Ziegler, 2020), possibly through the transcription factors Signal Transducer and Activator of Transcription (STAT) 1 and 2, as the authors suggest. The STAT family is comprised of seven transcription factors (STAT1, 2, 3, 4, 5A, 5B and 6) that are activated by type I and II cytokines through their respective receptors and the JAK/TYK2 family of tyrosine kinases (Stark and Darnell, 2012). Although each cytokine receptor has some preference for individual STAT members, it has become clear that any given cytokine can activate several, if not all, STAT members, which subsequently bind to a shared DNA motif, the gamma interferon activated sequence (GAS) (Hennighausen and Robinson, 2008). This permits individual genes to be activated by more than one cytokine through different receptors and several STAT family members.

While SARS-CoV-2 infection of lung epithelium is driving the disease, disturbances in other cell types (Qi et al., 2020), such as the olfactory system (Brann, 2020), have been observed. SARS-CoV-2 RNA has also been detected in breast milk of infected patients (Wu et al., 2020) suggesting that the virus can enter differentiated mammary alveolar cells and be vertically transmitted through breast feeding. Based on the overlapping activities of JAK-STAT components and their potential redundancy, it is likely that *ACE2* expression is activated by a wide range of cytokines through STATs 1, 2, 3 and 5. This has profound implications for strategies to mitigate ACE2 levels. Interfering with individual STATs will result in the compensational recruitment of other STAT members to cytokine receptors (Cui et al., 2007) with all its transcriptional consequences (Hennighausen and Robinson, 2008; Shin et al., 2016).

## Results and Discussion

*Ace2* mRNA levels vary widely between cell types, with high expression detected in lactating mammary and intestinal tissues (Figure 1A-B) and Type II Pneumocytes (Ziegler, 2020). To explore the possibility that *Ace2* gene expression in SARS-CoV-2 target cells is regulated not only by interferons but also by a range of cytokines through the family of STAT transcription factors, we mined available scRNA-seq data (Ziegler, 2020) (Table 1). Interferon receptors (IFNAR) and its downstream mediators JAK1, JAK2, TYK2 as well as STATs 1, 3 and 5 are highly expressed, thus supporting the mechanism of *ACE2* induction by IFN-α/β and IFN-γ. STAT1 levels increase sharply in cells treated with IFNs, supporting the notion of an autoregulatory loop (Yuasa and Hijikata, 2016). Moreover, these expression data point to the presence of functional STAT3 and STAT5 signaling cascades. Interleukin receptors, such as IL-7R, that are dependent on the common gamma chain (IL2RG), JAK1 and JAK3 are also highly expressed.

**Table 1.** mRNA levels of genes associated with the pan JAK-STAT pathway in primary human basal epithelial cells. scRNA-seq data were extracted from the study by Ziegler and colleagues(Ziegler, 2020). The human bronchial cell line (BEAS-2B) and airway basal cells from human donors had been exposed to interferons (IFNα2 and IFNγ) and cytokines (IL4 and IL17A). scRNA-seq libraries were generated with 15,000 cells. mRNA levels for genes in JAK/STAT signaling pathway were collected from the data and averages of independent biological replicates were normalized to the value of untreated group. Genes that were regulated more than 3-fold by interferons and cytokines are marked in red and highlighted colors.

| Cells | BEAS cell |  |  |  |  | Human1334 |  |  |  |  | Human2344 |  |  |  |  |
| --- | --- | --- | --- | --- | --- | --- | --- | --- | --- | --- | --- | --- | --- | --- | --- |
| N | 12 | 3 | 3 | 3 | 3 | 9 | 3 | 3 | 3 | 3 | 12 | 3 | 3 | 3 | 3 |
| drug (10ng/mL) | Untreated | IFNa2 | IFNg | IL4 | IL17A | Untreated | IFNa2 | IFNg | IL4 | IL17A | Untreated | IFNa2 | IFNg | IL4 | IL17A |
| ACE2 | 1.00 | 2.18 | 6.59 | 0.00 | 0.00 | 1.00 | 17.53 | 8.40 | 0.96 | 1.04 | 1.00 | 5.69 | 3.42 | 2.30 | 0.58 |
| TMPRSS2 | 1.00 | 0.19 | 0.59 | 0.30 | 0.51 | 1.00 | 0.52 | 0.53 | 2.42 | 1.03 | 1.00 | 0.74 | 0.71 | 1.38 | 1.49 |
| IFNAR1 | 1.00 | 0.78 | 1.01 | 0.87 | 1.11 | 1.00 | 0.76 | 0.61 | 0.86 | 0.97 | 1.00 | 0.79 | 0.56 | 0.76 | 0.66 |
| IFNAR2 | 1.00 | 1.26 | 1.72 | 0.90 | 1.20 | 1.00 | 1.05 | 1.33 | 0.90 | 0.75 | 1.00 | 0.80 | 1.06 | 0.70 | 0.94 |
| IFNGR1 | 1.00 | 1.08 | 0.74 | 0.52 | 1.32 | 1.00 | 1.01 | 0.74 | 0.74 | 1.70 | 1.00 | 0.97 | 0.64 | 0.94 | 1.20 |
| IL4R | 1.00 | 1.36 | 1.41 | 1.23 | 1.22 | 1.00 | 0.86 | 0.87 | 0.78 | 1.19 | 1.00 | 0.86 | 1.31 | 1.29 | 1.07 |
| IL12RB1 | 1.00 | 0.69 | 0.54 | 0.26 | 0.47 | 1.00 | 2.30 | 2.74 | 2.05 | 3.01 | 1.00 | 1.56 | 2.55 | 3.81 | 2.11 |
| IL12RB2 | 1.00 | 0.00 | 1.47 | 1.36 | 0.62 | 1.00 | 0.37 | 1.99 | 1.10 | 1.11 | 1.00 | 3.30 | 4.49 | 5.57 | 0.00 |
| IL13RA1 | 1.00 | 1.06 | 1.04 | 0.50 | 1.05 | 1.00 | 1.59 | 1.32 | 1.30 | 2.39 | 1.00 | 1.95 | 1.87 | 1.11 | 2.10 |
| IL13RA2 | 1.00 | 0.00 | 0.00 | 33.6 | 14.14 | 1.00 | 1.73 | 1.78 | 5.79 | 1.69 | 1.00 | 1.02 | 1.14 | 3.04 | 1.95 |
| IL15RA | 1.00 | 1.23 | 3.00 | 1.02 | 1.03 | 1.00 | 3.37 | 7.94 | 0.79 | 0.88 | 1.00 | 3.21 | 8.89 | 1.12 | 0.87 |
| IL17RA | 1.00 | 1.56 | 0.90 | 2.73 | 1.60 | 1.00 | 0.40 | 2.01 | 2.17 | 1.27 | 1.00 | 0.78 | 0.80 | 0.46 | 0.29 |
| IL17RC | 1.00 | 1.31 | 1.61 | 0.71 | 1.35 | 1.00 | 1.44 | 1.66 | 1.13 | 1.25 | 1.00 | 1.60 | 1.15 | 1.39 | 1.44 |
| STAT1 | 1.00 | 3.28 | 4.77 | 0.52 | 0.78 | 1.00 | 19.48 | 10.73 | 1.05 | 1.00 | 1.00 | 30.08 | 17.39 | 1.22 | 1.41 |
| STAT2 | 1.00 | 1.89 | 1.94 | 0.53 | 0.68 | 1.00 | 5.78 | 3.25 | 0.92 | 1.06 | 1.00 | 8.96 | 4.96 | 1.15 | 1.10 |
| STAT3 | 1.00 | 1.11 | 1.17 | 0.50 | 0.78 | 1.00 | 1.11 | 1.14 | 0.71 | 1.02 | 1.00 | 1.30 | 1.65 | 1.32 | 1.35 |
| STAT4 | 1.00 | 1.13 | 0.53 | 0.07 | 1.12 | 1.00 | 1.20 | 1.03 | 0.76 | 1.00 | 1.00 | 0.75 | 0.52 | 1.26 | 2.61 |
| STAT5A | 1.00 | 1.17 | 0.86 | 0.22 | 0.54 | 1.00 | 1.65 | 4.90 | 0.70 | 1.88 | 1.00 | 3.97 | 2.91 | 0.89 | 1.23 |
| STAT5B | 1.00 | 0.79 | 0.69 | 0.38 | 0.63 | 1.00 | 2.61 | 2.26 | 1.52 | 2.08 | 1.00 | 1.37 | 2.51 | 2.59 | 1.48 |
| STAT6 | 1.00 | 0.97 | 0.88 | 0.39 | 0.53 | 1.00 | 1.18 | 0.67 | 0.83 | 1.04 | 1.00 | 1.13 | 0.93 | 1.27 | 1.26 |
| JAK1 | 1.00 | 1.06 | 1.07 | 1.55 | 1.40 | 1.00 | 1.22 | 1.54 | 1.13 | 1.21 | 1.00 | 1.22 | 0.94 | 0.85 | 1.02 |
| JAK2 | 1.00 | 1.73 | 3.62 | 0.68 | 1.28 | 1.00 | 2.11 | 3.16 | 1.05 | 1.24 | 1.00 | 4.11 | 2.86 | 1.28 | 0.59 |
| JAK3 | 1.00 | 1.00 | 0.74 | 0.53 | 0.68 | 1.00 | 2.01 | 2.31 | 0.83 | 1.63 | 1.00 | 1.76 | 2.72 | 2.09 | 1.36 |
| TYK2 | 1.00 | 0.88 | 0.94 | 0.64 | 1.06 | 1.00 | 1.00 | 0.79 | 1.09 | 1.27 | 1.00 | 1.01 | 1.12 | 1.37 | 1.32 |
| LCN2 | 1.00 | 1.13 | 0.96 | 0.93 | 2.73 | 1.00 | 0.76 | 0.61 | 0.58 | 1.72 | 1.00 | 1.39 | 1.09 | 1.02 | 2.40 |
| FABP5 | 1.00 | 1.12 | 1.10 | 1.65 | 1.16 | 1.00 | 1.30 | 1.17 | 0.96 | 0.96 | 1.00 | 1.55 | 1.00 | 0.47 | 0.57 |
| AQP3 | 1.00 | 0.89 | 0.86 | 0.47 | 0.70 | 1.00 | 0.85 | 0.85 | 0.72 | 0.85 | 1.00 | 1.01 | 0.64 | 1.26 | 0.97 |
| SLC25A1 | 1.00 | 1.76 | 2.87 | 2.05 | 3.07 | 1.00 | 1.12 | 1.72 | 1.93 | 2.06 | 1.00 | 1.76 | 1.81 | 3.02 | 2.62 |
| SOCS2 | 1.00 | 1.43 | 1.13 | 0.65 | 1.47 | 1.00 | 1.21 | 0.45 | 0.96 | 0.90 | 1.00 | 0.64 | 0.69 | 2.64 | 2.18 |
| SOCS3 | 1.00 | 1.86 | 2.78 | 0.63 | 1.90 | 1.00 | 0.86 | 2.94 | 1.18 | 2.32 | 1.00 | 0.95 | 8.43 | 3.65 | 1.94 |
| BCL6 | 1.00 | 0.77 | 0.88 | 0.80 | 1.09 | 1.00 | 0.76 | 1.68 | 0.84 | 0.80 | 1.00 | 0.99 | 0.88 | 1.02 | 0.66 |
| CISH | 1.00 | 1.34 | 1.23 | 9.76 | 1.34 | 1.00 | 0.39 | 0.81 | 4.72 | 0.94 | 1.00 | 0.81 | 1.23 | 8.07 | 1.05 |

**Figure 1.**
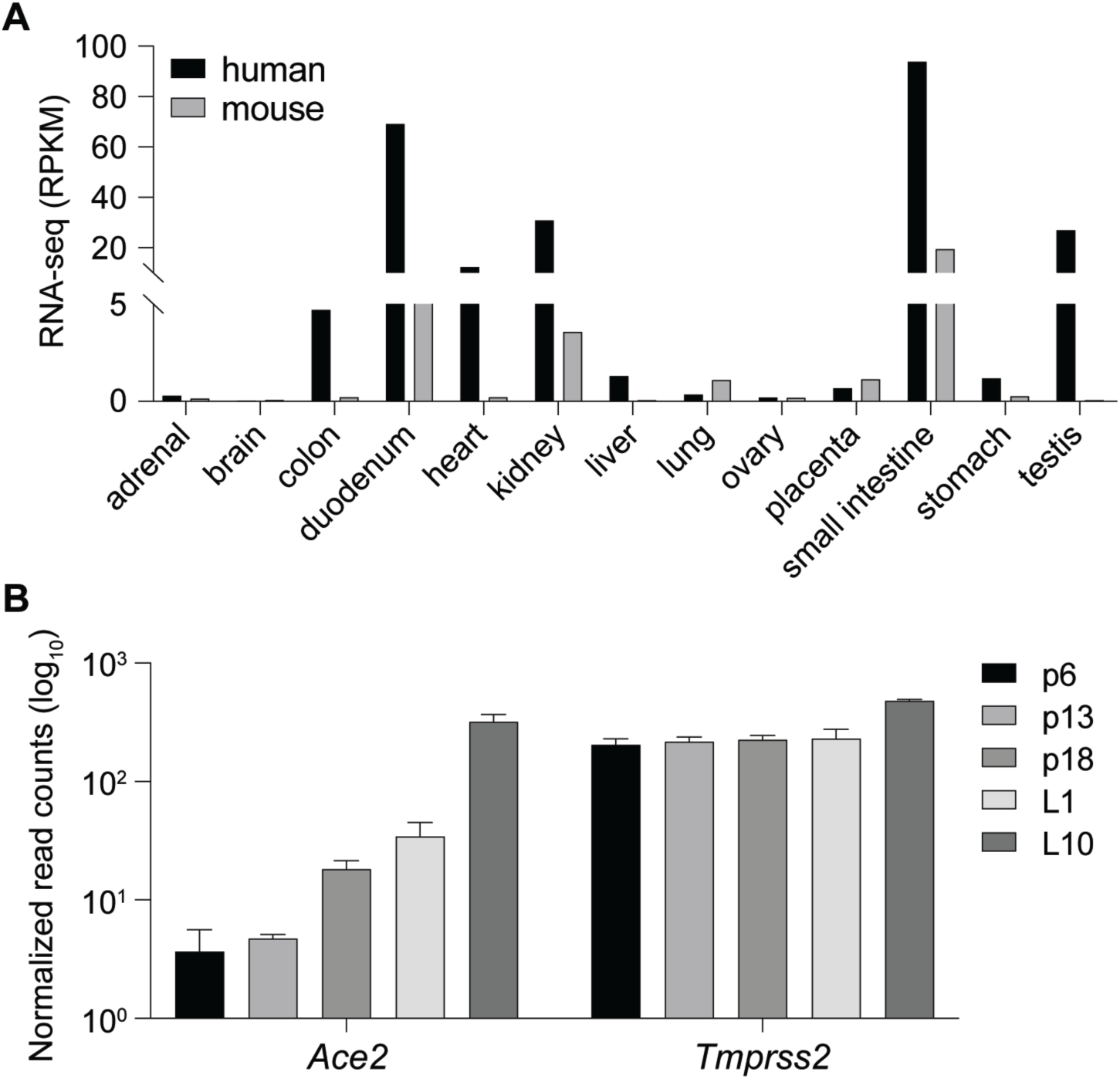
Ace2 is activated during lactation. **(A)** RNA-seq data from ENCODE demonstrate the presence of *ACE2* in several human and mouse tissues. **(B)** *Ace2* and *Tmprss2* mRNA levels in mouse mammary tissue at different stages of pregnancy and lactation were measured by RNA-seq. Day 6 of pregnancy (p6), p13, p18 and days 1 of lactation (L1) and L10.

The presence of a wide range of cytokine receptors, JAKs and STATs, suggests that *Ace2* might be activated by a broad selection of extracellular cues and most cytokines, including growth hormone and prolactin. We have tested this premise and explored whether *Ace2* is activated in mouse mammary tissue through STAT transcription factors. Gene expression in mammary epithelium during pregnancy and lactation is activated by prolactin through STAT5 (Liu et al., 1997). We observed an approximately 100-fold increase of *Ace2* mRNA during pregnancy and lactation (Figure 1B), which coincided with the establishment of a putative enhancer (Figure 2A). *Tmprss2* mRNA levels were similar throughout pregnancy and lactation (Figure 1B), suggesting that its expression is not under overt control of the JAK/STAT pathway. STAT5 was recruited to two distinct GAS (bona fide STAT binding motifs) in the candidate enhancer and co-occupancy of the glucocorticoid receptor (GR), nuclear factor 1 B (NFIB) and mediator complex subunit 1 (MED1) is likely not through their individual recognition motifs but through contacting STAT5. The presence of H3K4me1 enhancer marks, H3K27ac marks and RNA polymerase II (Pol II) occupancy further supports the validity of this regulatory region. Of note, no STAT3 occupancy was observed, suggesting a predominance of STAT5. In contrast to mammary tissue, limited STAT5 binding was observed in liver and no STAT5 and STAT3 binding was observed in kidney tissue (Figure 2B). The putative autoregulatory enhancer in the *Stat1* gene served as a positive control for STAT binding (Figure 2C).

**Figure 2.**
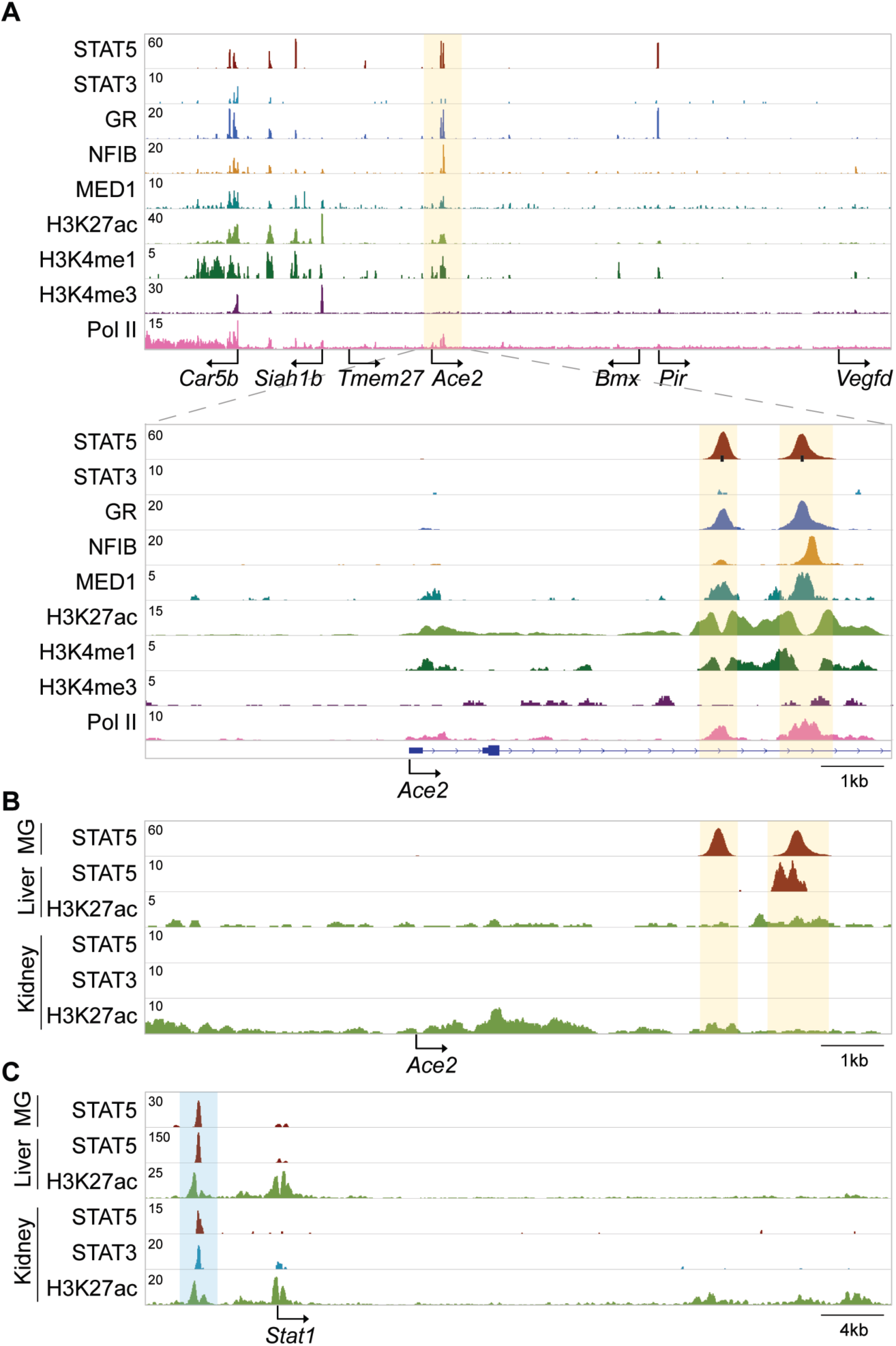
Establishment of a candidate *Ace2* enhancer during lactation. **(A)** ChIP-seq data for STAT5, STAT3, GR, NFIB, MED1 and histone markers H3K27ac and H3K4me3 provided structural information of the locus including the *Ace2* gene in day ten lactating mammary tissue. Solid arrows indicate the orientation of genes. The black bars indicate GAS motifs (STAT binding sites). The orange shades the candidate regulatory elements. **(B-C)** ChIP-seq profiles showed STAT binding at the candidate *Ace2* enhancer and the putative *Stat1* autoregulatory enhancer in mouse liver and kidney. The orange and blue shades indicate putative regulatory elements in each locus.

Our study demonstrates the presence of pan JAK/STAT components in Type II Pneumocytes, suggesting that *ACE2* is not only activated by IFN-α/β and IFN-γ but also by other cytokines. Moreover, we demonstrate an ∼100-fold increase of *Ace2* expression by pregnancy and lactation hormones in mouse mammary tissue. Future inquiries aimed at understanding the mechanism of *ACE2* gene regulation in potential SARS-CoV-2 target cells need to address the pan JAK-STAT pathway as well as steroid hormones, which might explain some of the sex differences seen in Covid-19 morbidity and mortality. Such investigations would need to include experimental approaches that comprehensively interrogate regulatory elements controlling *ACE2* expression *in vivo* in human tissues, both in males and females at different ages. Since underlying preexisting conditions, such as obesity, diabetes and high blood pressure, can affect the severity and progression of Covid-19, it would prudent to take this into account when analyzing the control of *Ace2* regulation.

## Materials and Methods

### Chromatin immunoprecipitation sequencing (ChIP-seq) analysis

Quality filtering and alignment of the raw reads was done using Trimmomatic (Bolger et al., 2014) (version 0.36) and Bowtie (Langmead et al., 2009) (version 1.1.2), with the parameter ‘-m 1’ to keep only uniquely mapped reads, using the reference genome mm10. Picard tools (Broad Institute. Picard, http://broadinstitute.github.io/picard/. 2016) was used to remove duplicates and subsequently, Homer (Heinz et al., 2010) (version 4.8.2) and deepTools (Ramirez et al., 2016) (version 3.1.3) software was applied to generate bedGraph files, seperately. Integrative Genomics Viewer (Thorvaldsdottir et al., 2013) (version 2.3.81) was used for visualization. Coverage plots were generated using Homer (Heinz et al., 2010) software with the bedGraph from deepTools as input. R and the packages dplyr (https://CRAN.R-project.org/package=dplyr) and ggplot2 (Love et al., 2014) were used for visualization. Each ChIP-seq experiment was conducted for two replicates. Sequence read numbers were calculated using Samtools (Masella et al., 2016) software with sorted bam files. The correlation between the ChIP-seq replicates was computed using deepTools using Spearman correlation.

### RNA-seq analysis

RNA-seq reads were analyzed using Trimmomatic (Bolger et al., 2014) (version 0.36) to check read quality (with following parameters: LEADING: 3, TRAILING: 3, SLIDINGWINDOW: 4:20, MINLEN: 36). The alignment was performed in Bowtie aligner (Langmead et al., 2009) (version 1.1.2) using paired end mode.

### Data availability

RNA-seq data from human and mouse tissues shown in Figure 1a were obtained from ENCODE. RNA-seq data shown in Fig. 1b and ChIP-seq data shown in Figure 2 were generated in our lab and deposited in the Gene Expression Omnibus (GEO) and ENCODE. ChIP-seq and RNA-seq data of mouse lactating tissue were obtained under GSE115370, GSE121438, GSE114294 and GSE127139. RNA-seq data of human bronchial cell line (BEAS-2B) and airway basal cells from human donors treated with IFNα2, IFNγ, IL4 or IL17A were obtained from GSE148829.

## Acknowledgments

This work was supported by the Intramural Research Program (IRP) of the National Institute of Diabetes and Digestive and Kidney Diseases (NIDDK) and utilized the computational resources of the NIH HPC Biowulf cluster (http://hpc.nih.gov).

## Author contributions

Conceptualization, H.K.L. and L.H.; Investigation, H.K.L.; Writing – Original Draft, L.H.; Writing – Review & Editing, H.K.L. and L.H.; Visualization, H.K.L; Supervision, H.K.L. and L.H.; Funding Acquisition, H.K.L. and L.H.

